# Habits are negatively regulated by HDAC3 in the dorsal striatum

**DOI:** 10.1101/153734

**Authors:** Melissa Malvaez, Venuz Y. Greenfield, Dina P. Matheos, Nicolas A. Angelillis, Michael D. Murphy, Pamela J. Kennedy, Marcelo A. Wood, Kate. M. Wassum

## Abstract

Optimal behavior results from a balance of control between two strategies, one cognitive/goal-directed and one habitual, which rely on the anatomically distinct dorsomedial (DMS) and dorsolateral (DLS) striatum, respectively. The transcriptional regulatory mechanisms required to learn and transition between these strategies are unknown. Here we identified a critical negative regulator of habit learning. Histone deacetylase (HDAC) inhibition following instrumental conditioning accelerated habitual control of behavior. HDAC3, a transcriptional repressor, was removed from the promoters of learning-related genes in the dorsal striatum as habits formed with overtraining and with post-training HDAC inhibition. Decreasing HDAC3 function in the DLS accelerated habit formation, while DLS HDAC3 overexpression prevented habit. HDAC3 activity in the DMS was also found to constrain habit formation. These results challenge the strict dissociation between DMS and DLS function in goal-directed v. habitual behavioral control and identify dorsal striatal HDAC3 as a critical molecular substrate of the transition to habit.

---

Decades of research have shown that humans and animals rely on two distinct strategies for reward seeking and decision making, one cognitive and one habitual (Dickinson, 1985; Dolan and Dayan, 2013). The deliberative, goal-directed strategy requires prospective evaluation of potential actions and their anticipated consequences, via a learned association between these variables, and, therefore supports behavior that can be readily adapted when circumstances change (Adams and Dickinson, 1981). Repetition of successful actions promotes less cognitively taxing habits, in which behavior is more automatically triggered by antecedent stimuli (Adams, 1982). The balance between these two forms of learning allows for adaptive and efficient behavior, but when it is disrupted can lead to the symptoms that underlie myriad psychiatric diseases (Gillan et al., 2016a; Voon et al., 2015).

Goal-directed and habit strategies have been demonstrated to rely on largely distinct cortico-basal ganglia circuits centered on the dorsomedial (DMS) (Corbit and Janak, 2010; Liljeholm et al., 2011; Yin et al., 2005a; Yin et al., 2005b) and dorsolateral (DLS) (Tricomi et al., 2009; Yin et al., 2004) striatum, respectively. Although gene transcription has been implicated in these persistent memory processes (Brightwell et al., 2008; Colombo et al., 2003; Hernandez et al., 2016; Hernandez et al., 2006; Pittenger et al., 2006; Shiflett et al., 2010), the molecular mechanisms that regulate transcriptional events that support the acquisition of and transition between these forms of behavioral control are unknown. Post-translational modifications of core histone proteins alter accessibility to DNA for transcriptional machinery to coordinate gene expression, and have been implicated in the regulation of neuronal plasticity and memory in several brain systems (Day and Sweatt, 2011; Kouzarides, 2007; Peixoto and Abel, 2013), and, thus, could regulate goal-directed and/or habit learning processes. But the function of such mechanisms in the dorsal striatum is not known. Histone deacetylases (HDACs) are a particularly interesting target because they are removed from gene promoters after salient behavioral events, allowing for histone acetylation, which in turn makes DNA accessible for transcriptional processes supporting the long-lasting changes in neuronal function that ultimately give rise to learning (Borrelli et al., 2008; McQuown and Wood, 2011; Vogel-Ciernia and Wood, 2012). Therefore, we evaluated the role of HDACs in instrumental action-reward learning using procedures that diagnose behavioral strategy combined with pharmacological and viral-mediated manipulation of HDAC function.

## RESULTS

### Effect of post-training HDAC inhibition on instrumental learning and behavioral strategy

Few studies have investigated the molecular mechanisms supporting the shift from goal-directed to habitual control of behavior. We first examined the function of HDACs in this process by systemically inhibiting HDAC activity following instrumental conditioning and then probing behavioral strategy off drug. Rats were trained to press a lever to earn delivery of a food-pellet reward on a random-interval 30-s schedule of reinforcement. Two groups of subjects were given either limited or intermediate training, known to preserve goal-directed control, while a third group was given extended training known to allow habits to dominate (Adams, 1982). Rats were administered the nonspecific class I HDAC inhibitor sodium butyrate (Kilgore et al., 2010) (NaBut, 1.0 g/kg, i.p.) or sterile water vehicle immediately after each training session to examine the function of HDACs in consolidation of the learning underlying instrumental performance. All rats acquired the instrumental behavior; both vehicle- and NaBut-treated rats increased their lever-press rate across training days (Fig. 1A-C; main effect of Training day: Limited, *F*_3,60_=92.59, *P*<0.0001; Intermediate, *F*_4,80_=50.69, *P*<0.0001; Extended, *F*_6,96_=18.64, *P*<0.0001), with the NaBut group plateauing at a lower rate than vehicle controls with limited (Drug x Day: *F*_3,60_=3.63, *P*=0.02), or intermediate (*F*_4,80_=5.49, *P*<0.001), but not extended (*F*_6,96_=0.42, *P*=0.86) training.

**Figure 1.**
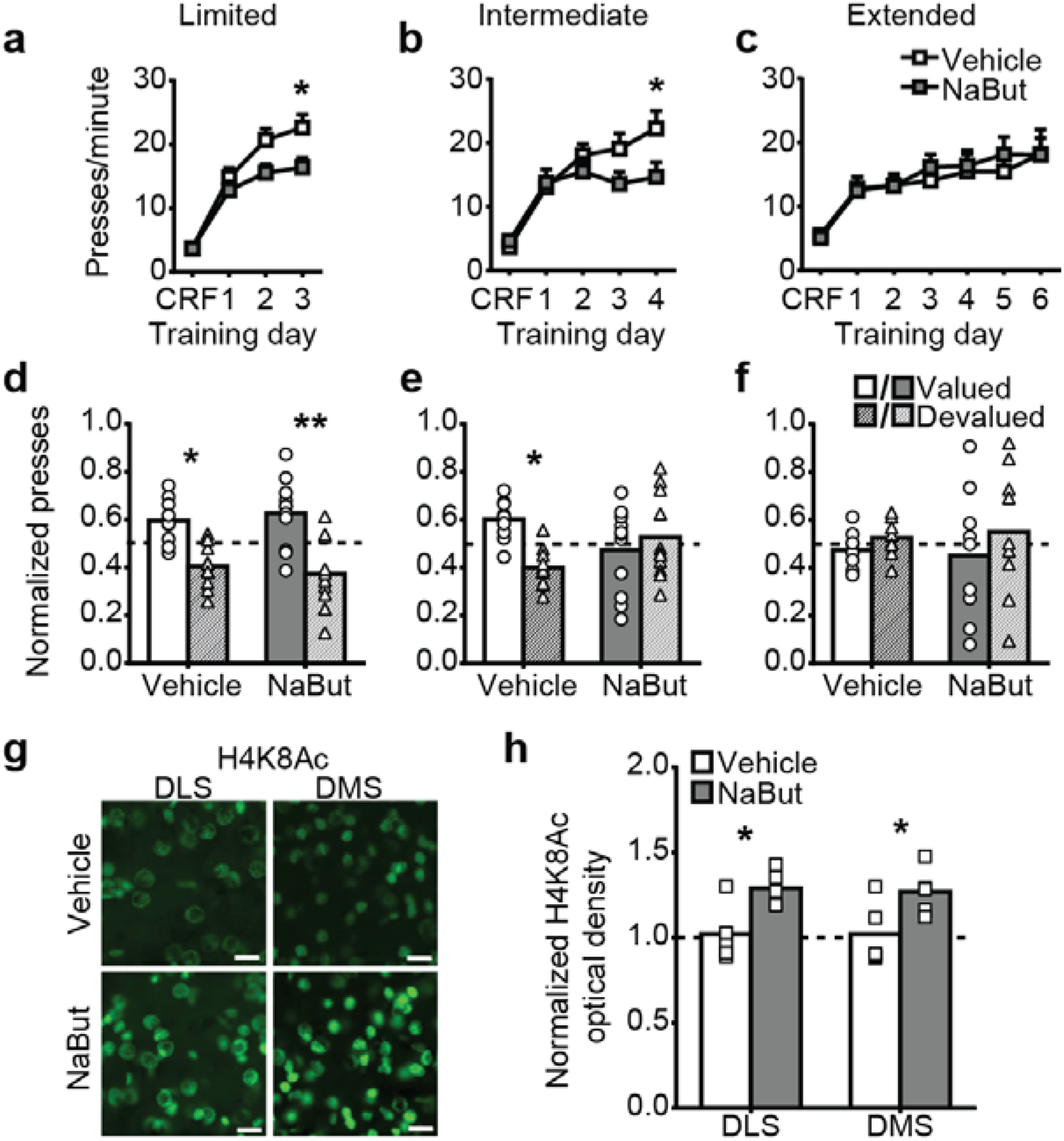
Effect of post-training HDAC inhibition on instrumental learning, behavioral strategy, and acetylation at histone H4. (**a-c**) Instrumental (lever press→reward) training performance for rats given limited (**a**; *N*=11/group), intermediate **(b**; *N*=11-12/group), or extended (**c**; *N*=9/group) instrumental training. (data presented as mean + s.e.m). (**d-f**) Lever presses during the subsequent devaluation tests normalized to total presses across both tests for the Valued [Valued state presses/(Valued + Devalued state presses)] and Devalued [Devalued state presses/(Valued + Devalued state presses)] states for rats that received limited (**d**), intermediate (**e**), or extended (**f**) training. Dashed line indicates point of equal responding between tests. (data presented as mean + scatter). (**g, h**) Representative immunofluorescent image (**g**) and quantification (**h**) of acetylation of H4K8 (H4K8Ac; *N*=4-6/condition) 1 hr following instrumental training/drug treatment. Data normalized to vehicle control (dashed line). \**P*<0.05; \*\**P*<0.01.

Behavioral strategy cannot be determined from simple lever-press performance. To identify goal-directed v. habitual control, rats were given a brief, 5-min, drug-free, outcome-specific devaluation test (Fig. 1D-F). Non-reinforced lever pressing was assessed following sensory-specific satiation (1 hr pre-feeding) on the food pellet earned by lever pressing (Devalued state) and compared to pressing after satiation on an alternate food pellet that had been non-contingently provided daily outside of the training context (Valued state). Each rat was tested under both conditions and data were normalized across tests. If subjects are using a goal-directed strategy and, therefore, considering the consequences of their lever-press actions, they should downshift their pressing in the devalued state (‘sensitivity to devaluation’). Habits are marked by insensitivity to outcome devaluation. As expected, in vehicle-treated subjects we observed a reduction in lever pressing in the Devalued state relative to the Valued state in the limited training group (Fig. 1D; Devaluation: *F*_1,20_=19.68, *P*=0.003) and a failure to reduce responding following devaluation in the extended training group (Fig. 1F; *F*_1,16_=0.62, *P*>0.999). This was not altered by post-training HDAC treatment (Drug: Limited, *F*_1,20_=0.00, *P*>0.999; Extended, *F*_1,16_=1.88, *P*=0.19; Drug x Devaluation: Limited, *F*_1,20_=0.41, *P*=0.53; Extended, *F*_1,16_=0.06, *P*=0.80), suggesting that HDAC inhibition neither disrupted initial goal-directed learning, nor prevented the normal development of habits associated with overtraining.

Post-training HDAC inhibition did, however, accelerate the rate at which the habit strategy came to dominate behavioral control. With intermediate training, NaBut-treated animals became insensitive to devaluation of the earned reward, failing to reduce responding in the Devalued v. Valued states, while lever pressing in vehicle-treated subjects remained sensitive to devaluation (Fig. 1E; Devaluation: *F*_1,20_=1.68, *P*=0.21; Drug: *F*_1,20_=0.74, *P*=0.40; Drug x Devaluation: *F*_1,20_=4.85, *P*=0.04). In separate groups, we also found that post-training HDAC inhibition accelerated habit formation when behavioral strategy was assayed by reversal of the action-outcome contingency (Fig. S1) and occurred even when rats were trained on a random-ratio reinforcement schedule (Fig. S2) known to preserve goal-directed control of action (Dickinson, 1985). Importantly, this effect did not manifest if NaBut was administered well outside the memory-consolidation window (Fig. S3). Together, these data reveal that inhibition of class I HDACs following instrumental conditioning facilitates transition to habitual control of behavior, allowing habits to dominate at a point in training that behavior would normally remain primarily goal-directed.

### Effect of post-training HDAC inhibition on dorsal striatal activity and histone acetylation

HDACs constrain histone acetylation levels by removing acetyl groups from histone tails (Kouzarides, 2007), therefore, HDAC inhibitor treatment should permit increased histone acetylation (Peixoto and Abel, 2013). We next used immunofluorescence to determine the brain region-specific effects of peripheral NaBut treatment, and hypothesized histone acetylation would be increased in the DLS, a region critical for habit formation (Yin et al., 2004). We focused on histone H4 lysine 8 (H4K8Ac), a known target of the most highly-expressed class I HDAC in the brain, HDAC3 (Broide et al., 2007; McQuown et al., 2011), and implicated in memory formation (Kwapis et al., 2017; McQuown et al., 2011). 1 hr after the last intermediate training session and drug delivery, H4K8Ac was significantly higher in the DLS (*t*_8_=3.08, *P*=0.02) of NaBut-treated subjects compared to controls (Fig. 1 G-H). H4K8Ac was also elevated in the DMS (*t*_8_=2.41, *P*=0.04), but there were no significant differences in any other regions examined (see Table S1).

To determine whether post-training HDAC inhibition altered activity in the dorsal striatum, we measured expression of the immediately early gene (IEG) neuronal activity markers *c-fos, Egr1*, and *FosB* implicated in memory processes, including instrumental acquisition conditioning (Colombo et al., 2003; Hernandez et al., 2006). IEG expression was significantly increased in the dorsal striatum 1 hr after instrumental training, with no significant differences between training or drug groups (Fig. S4), indicating that both the DMS and DLS are engaged during instrumental conditioning and that this is not significantly altered by post-training HDAC inhibition.

### Effect of training and post-training HDAC inhibition on HDAC3 occupancy at learning-related gene promoters and gene expression

In general, elevated histone acetylation is associated with a transcriptionally-permissive state (Kouzarides, 2007), such that HDAC activity typically represses gene expression. That post-training HDAC inhibition accelerated the transition to habit, suggests that HDACs might normally be engaged in restraining the gene expression underlying habit formation. If this is the case, then HDAC occupancy at the promoters of key learning-related genes might be relatively high with intermediate levels of instrumental training to prevent dominance by the habit strategy, but removed when conditions are ripe for habit to dominate, e.g., overtraining, or by HDAC inhibition which accelerates this form of learning. To test this, we used chromatin immunoprecipitation (ChIP) following the last training session to examine HDAC occupancy at the promoter regions of *Bdnf1*, *Nr4a1*, and *Nr4a2* (Fig 2A-C). These genes were selected because they have been shown to be regulated by histone acetylation, involved in long-term memory formation (Bredy et al., 2007; Intlekofer et al., 2013; Kwapis et al., 2017; Malvaez et al., 2013; McQuown et al., 2011), and are regulated by CREB (McNulty et al., 2012; Ou and Gean, 2007; Vecsey et al., 2007), a transcription factor implicated in habit-like behaviors (Brightwell et al., 2008; Colombo et al., 2003; Pittenger et al., 2006). We focused on the DLS, given its crucial role in habit formation and findings of training-related activity and NaBut regulation of H4K8Ac in this region, and HDAC3, the most highly expressed class I HDAC in the striatum (Broide et al., 2007).

**Figure 2.**
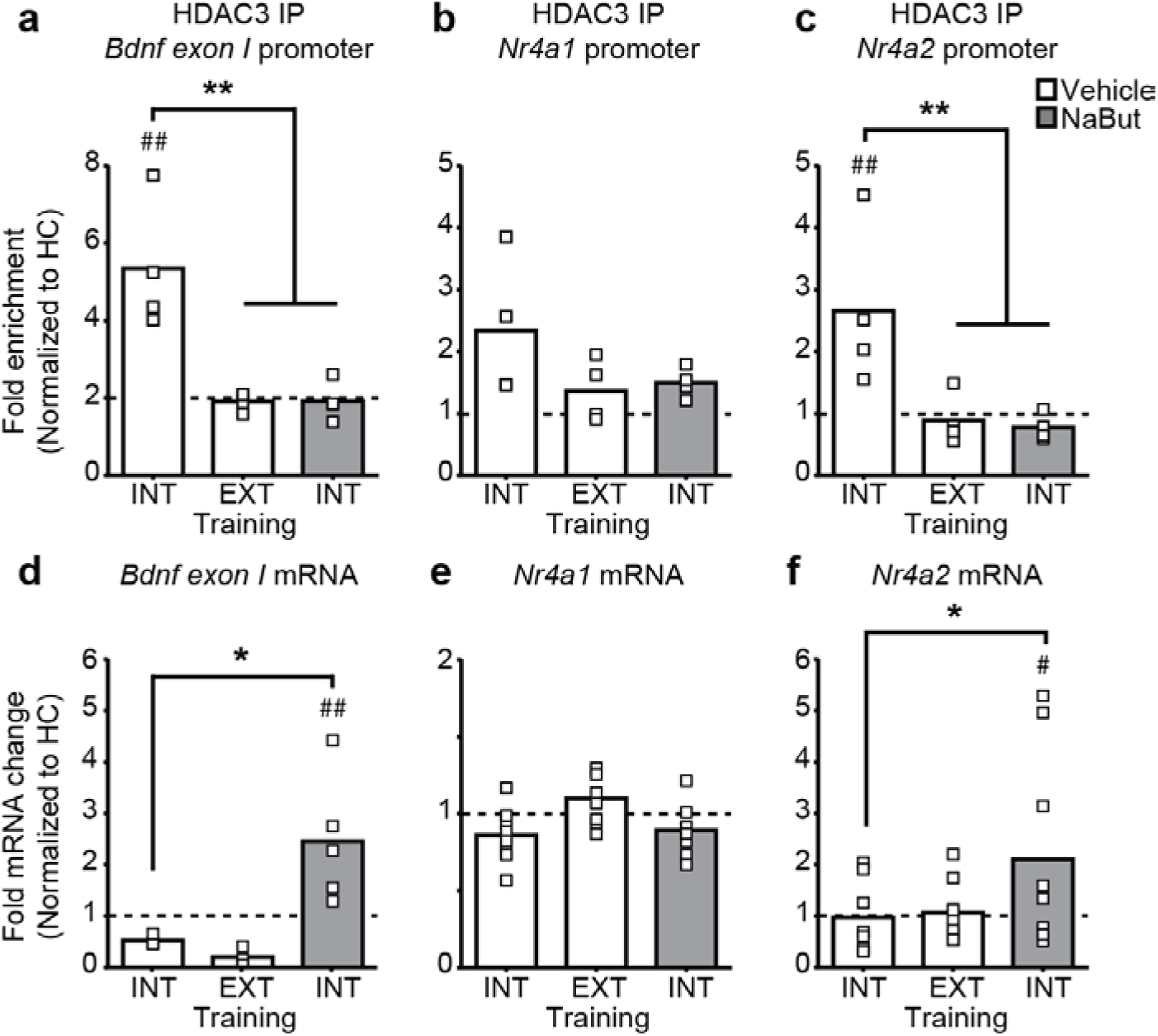
Effect of training and post-training HDAC inhibition on HDAC3 occupancy at learning-related gene promoters and gene expression in the dorsolateral striatum. (**a-c**) ChIP was performed with anti-HDAC3 followed by qPCR to identify HDAC3 binding to the *Bdnf1* (**a**), *Nr4a1* (**b**), *or Nr4a2* (**c**) promoters in the DLS following either intermediate (INT) or extended training (EXT) in vehicle-treated rats, or NaBut treatment post-intermediate training, relative to home cage control levels (dashed line). (**d-e**) mRNA expression of *Bdnf1* (**d**), *Nr4a1* (**e**), and *Nr4a2* (**f**) in the DLS. \**P*<0.05, \*\**P*<0.01, between groups; ^##^*P*<0.01 relative to homecage control.

Following intermediate training, when behavior was still goal-directed, HDAC3 occupancy at the *Bdnf exon 1* promoter was enriched relative to homecage controls (*P*<0.05), but returned to control levels with the extended training that promotes habits and was not elevated when intermediate training was followed by NaBut treatment (Fig. 2A; *F_3,12_*=18.53, *P*<0.001). A similar pattern was detected for HDAC3 occupancy at the promoter for *Nr4a2* (Fig. 2C; *F_3,12_*=5.74, *P*=0.01) and at *Nr4a1*, though was less robust in this case (Fig. 2B; *F_3,12_*=3.18, *P*=0.06). Expression of *Bdnf1* (Fig. 2D) and *Nr4a2* (Fig. 2F) was significantly elevated 1 hr following intermediate training with NaBut treatment relative to homecage controls (*BDNF*: t_13_=2.41, *P*=0.03; *Nr4a2*: t_31_=2.16, *P*=0.04) and the intermediate trained control group (*BDNF*: t_13_=3.02, *P*=0.01; *Nr4a2*: t_31_=2.14, *P*=0.04), but was not significantly different from controls following extended training. *Nr4a1* expression (Fig. 2E) was not significantly different from homecage controls in any condition.

These data suggest that HDAC3 occupancy at the *Bdnf1* and *Nr4a2* promoters is normally enriched when instrumental behavior is goal-directed and that HDAC3 becomes dissociated from these gene promoters as habits dominate behavioral control. Post-training HDAC inhibition accelerates both habit formation and removal of HDAC3 from the promoters of these key learning genes, and also induces *Bdnf1* and *Nr4a2* expression. Therefore, HDAC inhibition immediately following instrumental training caused a transcriptionally-permissive, hyperacetylated histone state, disengaged HDAC3 from specific gene promoters in the DLS, and facilitated behavioral control by the habit system.

### Effect of HDAC3 manipulation in dorsolateral striatum on habit formation

The data show that HDAC3 is disengaged in the DLS following extended instrumental training and with HDAC inhibition accelerated habit. This suggests that habits might be promoted by decreased HDAC3 activity in the DLS. We tested this hypothesis in two complementary ways. First by pharmacologically inhibiting HDAC3 activity specifically in the DLS after intermediate instrumental training and, second, by expressing a dominant negative HDAC3 point mutant (Kwapis et al., 2017) in the DLS to selectively disrupt HDAC3 enzymatic activity, without affecting protein-protein interactions (Lahm et al., 2007; Sun et al., 2013). In both cases, habit formation was potentiated, recapitulating the effects of systemic post-training HDAC inhibition. Post-training intra-DLS infusion of the selective HDAC3 inhibitor RGFP966 (Bieszczad et al., 2015; Malvaez et al., 2013; Rumbaugh et al., 2015) elevated H4K8Ac in the DLS (Fig. 3A-B; *t*_8_=3.15, *P*=0.014), but not the adjacent DMS (Fig. 3A; Normalized H4K8Ac optical density: Vehicle, 1.011±0.10; RGFP966, 1.00±0.27; *t*_3_=0.06, *P*=0.954) compared to vehicle-treated controls. This treatment did not alter the acquisition of instrumental lever-press behavior (Fig. 3C; Training day: *F*_4,68_=32.61, *P*<0.001; Drug: *F*_1,17_=0.95, *P*=0.34; Drug x Day: *F*_4,68_=2.32, *P*=0.07), but did render this behavior insensitive to devaluation of the earned reward under conditions in which intermediate trained controls remained sensitive (Fig. 3D; Devaluation: *F_1,17_*=10.83, *P*=0.004; Drug: *F*_1,17_=1.12, *P*=0.31; Drug x Devaluation: *F*_1,17_=7.25, *P*=0.02). Similarly, expressing a dominant negative point mutant of HDAC3 (AAV2/1-CMV-HDAC3Y298H-V5), in the DLS produced H4K8 hyperacetylation (Fig. 3E-F; *t*_10_=3.91, *P*=0.003) and insensitivity to outcome devaluation (Fig. 3H; Devaluation: *F*_1,14_=3.74 *P*=0.07; Virus: *F*_1,14_=0.47, *P*=0.506; Virus x Devaluation: *F*_1, 17_=6.19, *P*=0.03), without altering lever-pressing acquisition (Fig. 3G; Training day: *F*_4,56_=29.21, *P*<0.001; Virus: *F*_1,14_<0.01, *P*=0.99; Virus x Day: *F*_4,56_=0.56, *P*=0.69). These results demonstrate that blocking HDAC3 activity in the DLS promotes habitual control of instrumental behavior.

**Figure 3.**
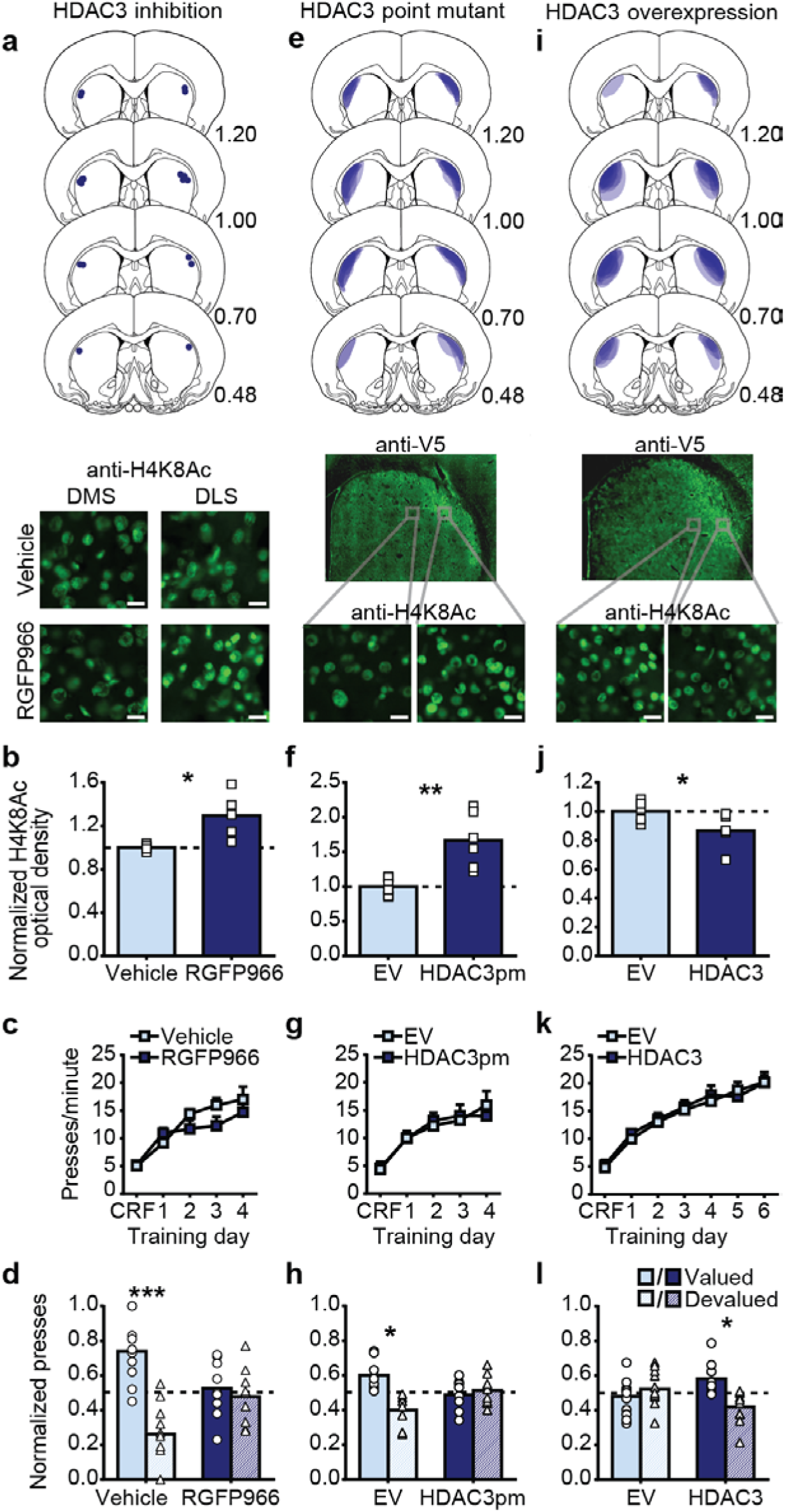
Effect of HDAC3 manipulation in dorsolateral striatum on habit formation. (**a**) *Top*, schematic representation of injector tips in the DLS. Numbers to the lower right of each section represent distance (mm) anterior to bregma. Coronal section drawings taken from (Paxinos and Watson, 1998). *Middle*, representative immunofluorescent images of H4K8Ac in DIVIS (*left)* and DLS (*right)* 1 hr following instrumental training/intra-DLS vehicle (*top*) or RGFP966 (*bottom*) infusion. (**e, i**) *Top*, schematic representation of HDAC3 point mutant (**e**), or HDAC3 (**i**) expression in the DLS for all subjects. *Middle*, representative immunofluorescent images of V5-tagged HDAC3 point mutant (**e**) or HDAC3 (**i**) expression in the DLS. *Bottom*, Representative immunofluorescent images of H4K8Ac in rats expressing the HDAC3 point mutant (**e**) or HDAC3 (**i**) either outside (*left*) or inside (*right*) of expression zone. (**b, f, j**) Quantification of H4K8Ac for rats receiving intra-DLS vehicle or RGFP966 infusion (**b**; *N*=5/group), intra-DLS empty vector (EV) or HDAC3 point mutant (HDAC3pm) (**f**; *N*=6/group), or intra-DLS empty vector or HDAC3 overexpression (**j**; *N*=5-6/group). (data presented as mean + scatter). (**c, g, k**) Instrumental training performance for rats given post-training intra-DLS vehicle or RGFP966 infusions (**c**; *N*=10-12/group), intra-DLS empty vector or HDAC3 point mutant (**g**; *N*=8/group), or intra-DLS empty vector or HDAC3 (**k**; *N*=11-13/group). (data presented as mean + s.e.m). (**d, h, l**) Normalized lever presses during the subsequent devaluation tests for rats given post-training intra-DLS vehicle or RGFP966 infusions (**d**), intra-DLS empty vector or HDAC3 point mutant (**h**), or intra-DLS empty vector or HDAC3 (**l**). \**P*<0.05; \*\**P*<0.01; \*\*\**P*<0.001.

The combined data (Fig. 2 and 3) strongly suggest that in the DLS HDAC3 might normally be engaged in restraining habits when sufficient action repetition has not yet occurred. If this is the case, then increasing HDAC3 activity in the DLS should slow or prevent habit formation. To test this, we overexpressed HDAC3 in the DLS (AAV2/1-CMV-HDAC3-V5), confirmed that this reduced H4K8Ac in the DLS (Fig. 3I-J; *t*_9_=2.29, *P*=0.048), and gave subjects extended instrumental training known to promote habit formation. All subjects similarly acquired the lever-press behavior (Fig. 3K; Training day: *F*_6,126_=81.44, *P*<0.001; Virus: *F*_1,21_=0.06, *P*=0.81; Virus x Day: *F*_6,126_=0.44, *P*=0.85). As expected, control subjects showed evidence of habits (insensitivity to devaluation), whereas subjects with HDAC3 overexpressed in the DLS were unable to form behavioral habits and continued to show sensitivity to devaluation (Fig. 3L; Devaluation: *F*_1,21_=2.24 *P*=0.15; Virus: *F*_1,21_=-8.17, *P*>0.999; Virus x Devaluation: *F*_1,21_=6.45, *P*=0.02), even after extensive overtraining (Fig. S5). Together, these data demonstrate that HDAC3 in the DLS is a critical negative regulator of habit formation.

### Effect of HDAC3 manipulation in dorsomedial striatum on habit formation

We found that systemic post-instrumental-training HDAC inhibition increased H4K8Ac in both the DLS and DMS. In order to further understand how altered regulation of gene transcription in these dissociable brain systems contributes to a transition in behavioral control we next asked whether manipulating HDAC3 in the DMS would modify the progression of instrumental strategy. Both post-training intra-DMS HDAC3 inhibition (RGFP966 infusion; Fig. 4A-B; DMS: *t*_7_=2.54, *P*=0.04; DLS: Vehicle, 1.00±0.02 normalized H4K8Ac optical density v. RGFP966 1.09±0.07, *t*_7_=1.30, *P*=0.23) and expression of the dominant negative HDAC3 point mutant in the DMS (Fig. 4E-F; *t*_7_=4.32, *P*=0.004) induced a hyperacetylated (elevated H4K8Ac) state that was restricted to the DMS. Neither treatment affected acquisition of the lever-press behavior with intermediate training (RGFP966: Fig. 4C; Training day: *F*_4,64_=45.38, *P*<0.001; Drug: *F*_1,16_=0.06, *P*=0.81; Drug x Day; *F*_4,64_=0.12, *P*=0.98; HDAC3pm: Fig. 4G; Day: *F*_4,84_=32.38, *P*<0.001; Virus: *F*_1,21_=0.19 *P*=0.67; Virus x Day: *F*_4,84_=0.44, *P*=0.78). To our surprise, given the canonical function of the DMS in goal-directed not habit learning (Yin et al., 2005a; Yin et al., 2005b), both intra-DMS RGFP966 (Fig. 4D; Devaluation: *F*_1,16_=1.16 *P*=0.298; Drug: *F*_1,16_=0.00, *P*>0.999; Drug x Devaluation: *F*_1, 16_=4.79, *P*=0.04) and DMS HDAC3 point mutant expression (Fig. 4H; Devaluation: *F*_1,21_=1.94, *P*=0.18; Virus: *F*_1,21_=17.61, *P*<0.001; Virus x Devaluation: *F*_1,21_=5.52, *P*=0.03) potentiated habit formation, as indicated by insensitivity to devaluation of the earned reward following intermediate training. Conversely, DMS HDAC3 overexpression reduced H4K8Ac (Fig. 4I and 3J; *t*_8_=3.27, *P*=0.01) did not alter lever pressing during extended training (Fig. 3K; Day: *F*_6,132_=61.69, *P*<0.001; Virus: *F*_1,22_=1.36, *P*=0.26; Virus x Day: *F*_6,132_=2.01, *P*=0.07), but prevented habit formation (Fig. 3L; Devaluation: *F*_1,22_=7.06, *P*=0.01; Virus: *F*_1,22_=2.68, *P*=0.12; Virus x Devaluation: *F*_1,22_=4.34, *P*=0.049), even with extensive overtraining (Fig. S6). These unexpected results demonstrate that, like in the DLS, HDAC3 activity in the DMS negatively regulates the transition to habit.

**Figure 4.**
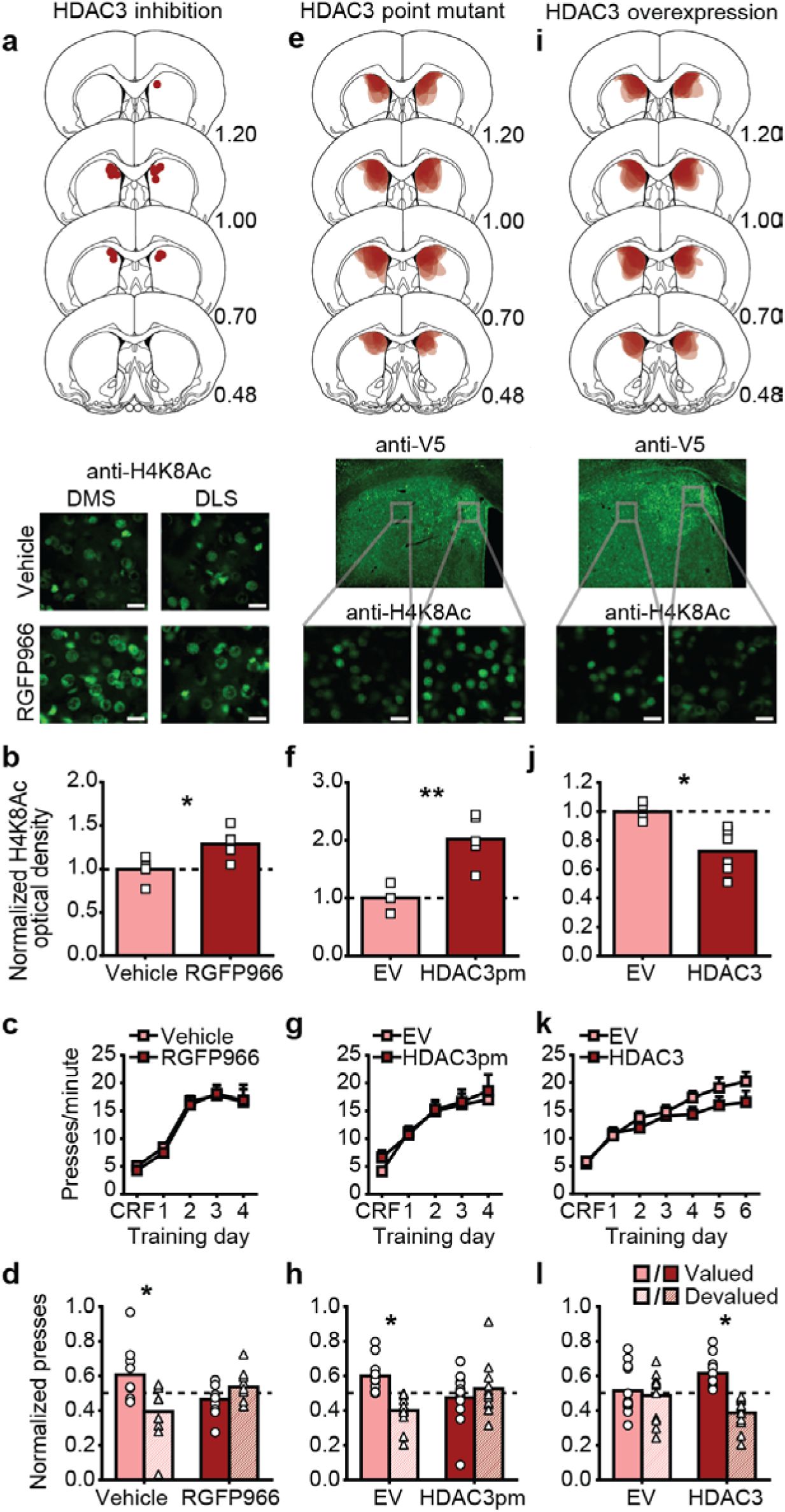
Effect of HDAC3 manipulation in dorsolateral striatum on habit formation. (**a**) *Top*, schematic representation of injector tips in the DMS. Numbers to the lower right of each section represent distance (mm) anterior to bregma. *Middle*, representative immunofluorescent images of H4K8Ac in DMS (*left)* and DLS (*right)* 1 hr following instrumental training/intra-DMS vehicle (*top*) or RGFP966 (*bottom*) infusion. (**e, i**) *Top*, schematic representation of HDAC3 point mutant (**e**), or HDAC3 (**i**) expression in the DMS for all subjects. *Middle*, representative immunofluorescent images of HDAC3 point mutant (**e**) or HDAC3 (**i**) expression in the DMS. *Bottom*, Representative immunofluorescent images of H4K8Ac in rats expressing the HDAC3 point mutant (**e**) or HDAC3 (**i**) either outside (*left*) or inside (*right*) of the expression zone. (**b, f, j**) Quantification of H4K8Ac for rats receiving intra-DMS vehicle or RGFP966 infusion (**b**; *N*=4-5/group), intra-DMS empty vector (EV) or HDAC3 point mutant (**f**; *N*=4-5/group), or intra-DMS empty vector or HDAC3 overexpression (**j**; *N*=4-6/group). (data presented as mean + scatter). (**c, g, k**) Instrumental training performance for rats given post-training intra-DMS vehicle or RGFP966 infusions (**c**; *N*=9/group), intra-DMS empty vector or HDAC3 point mutant (**g**; *N*=11-12/group), or intra-DMS empty vector or HDAC3 (**k**; *N*=12/group). (data presented as mean + s.e.m). (**d, h, l**) Normalized lever presses during the subsequent devaluation tests for rats given post-training intra-DMS vehicle or RGFP966 infusions (**d**), intra-DMS empty vector or HDAC3 point mutant (**h**), or intra-DMS empty vector or HDAC3 (**l**). \**P*<0.05; \*\**P*<0.01.

Given this surprising finding, we next examined whether HDAC3 is normally engaged in the DMS with instrumental training and as habits develop. Using ChIP in the DMS (Fig 5A-C), we found that, unlike in the DLS, HDAC3 occupancy at *Bdnf1* was not significantly altered by instrumental training or intermediate training followed by systemic NaBut treatment (Fig. 5A; *F_3,12_*=2.288, *P*=0.13). Although there was an overall effect of training/treatment group on HDAC3 occupancy at the *Nr4a2* promoter (Fig. 5C; *F_3,12_*=4.05, *P*=0.03), in no case was HDAC3 enrichment at this promoter significantly different from homecage controls (*P*>0.05). Instead, HDAC3 occupancy at the *Nr4a1* promoter (Fig. 5B; *F_3,12_*=7.76, *P*=0.004) was increased relative to homecage controls with intermediate training (*P*<0.01) and returned to control levels following extended training and was not elevated following intermediate training with NaBut treatment. Follow up mRNA analysis revealed no significant changes in expression of these genes (Fig. 5D-F; *Bdnf1, F_3,16_*=1.57, *P*=0.24; *Nr4a1, F_3,25_*=1.87, *P*=0.16; *Nr4a2, F_3,30_*=2.49, *P*=0.079). These results suggest that HDAC3 activity in the DMS and DLS is differentially regulated by instrumental training as habits form.

**Figure 5.**
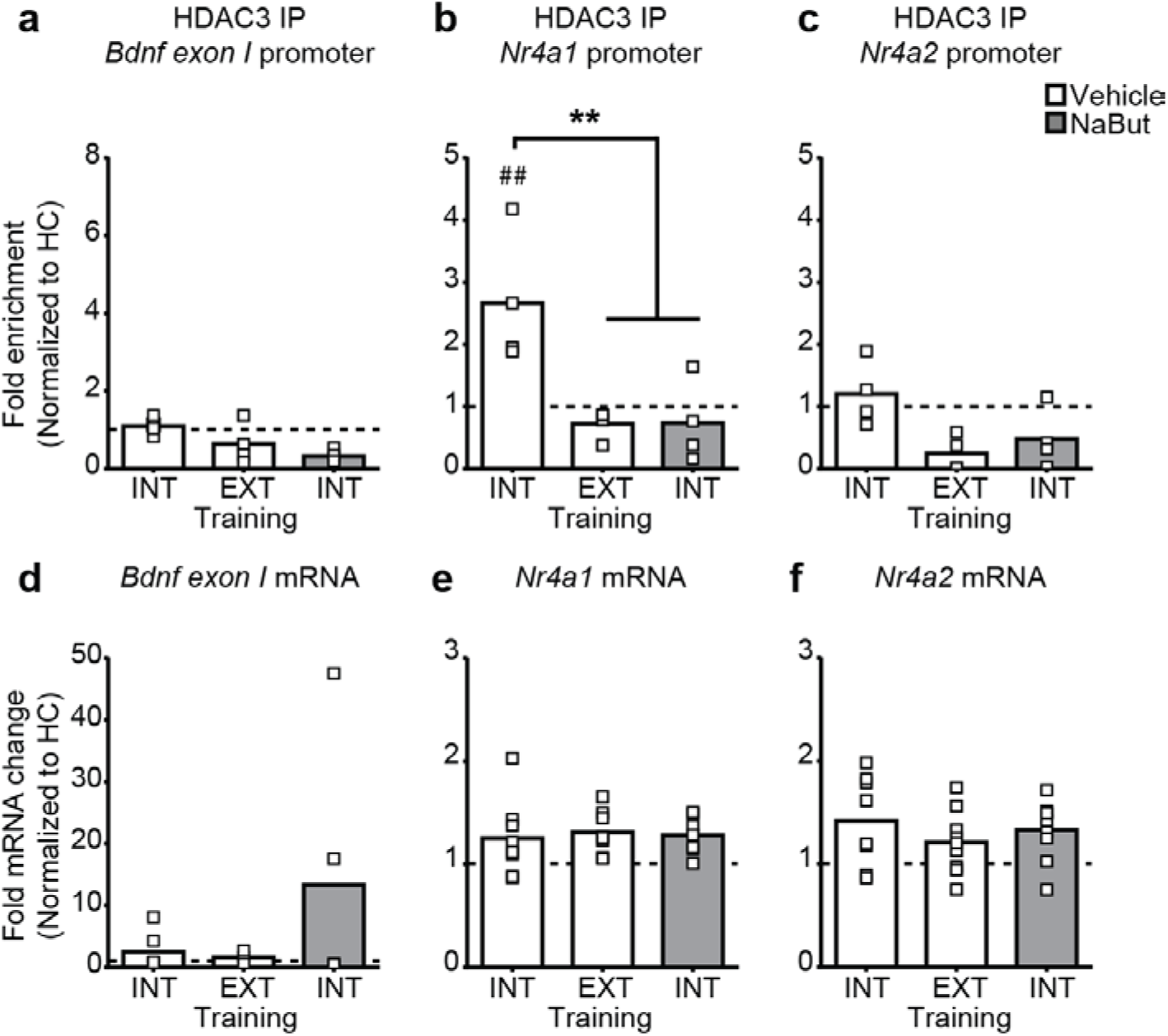
Effect of training and post-training HDAC inhibition on HDAC3 occupancy at learning-related gene promoters and gene expression in the dorsomedial striatum. (**a-c**) ChIP was performed with anti-HDAC3 followed by qPCR to identify HDAC3 binding to the *Bdnf1* (**a**), *Nr4a1* (**b**), *or Nr4a2* (**c**) promoters in the DMS following either intermediate training (INT) or extended training (EXT) in vehicle-treated rats, or NaBut treatment post-intermediate training, relative to home cage control levels (dashed line). (**d-f**) mRNA expression of *Bdnf1* (**d**), *Nr4a1* (**e**), and *Nr4a2* (**f**) in the DMS. \*\**P*<0.01, between groups; ^##^*P*<0.01 relative to homecage control.

## DISCUSSION

Reward seeking and decision making are controlled by a delicate balance between two control systems, one reflective, involving prospective consideration of learned action consequences, and one reflexive, allowing common behaviors to be automatically triggered by antecedent events on the basis of their past success. Here we identified a critical molecular regulator of the transition of behavioral control to the latter, habit, strategy: dorsal striatal HDAC3. Systemic HDAC inhibition during instrumental acquisition increases histone acetylation in the dorsal striatum and accelerates habitual control of behavior. HDAC3 occupancy at specific learning-related gene promoters in the dorsal striatum is reduced when habits form with overtraining and this is mimicked by HDAC inhibition. HDAC3 activity constrains habit learning not only in the DLS, previously implicated in habit (Yin et al., 2004), but also in the DMS, which has been canonically ascribed a role in the opposing, goal-directed, strategy (Yin et al., 2005a; Yin et al., 2005b).

HDAC3 in the dorsal striatum functions as a negative regulator of habit, as evidenced by its disruption accelerating the rate at which behavioral control transitioned to dominance by the habit system and its overexpression in the dorsal striatum preventing subjects from forming habits under conditions that would normally promote them to do so. The temporally-restricted effect of post-training HDAC inhibition suggests that HDAC3 negatively regulates the consolidation of habit memories, similar to previous findings of HDAC regulation of memory processes in other brain regions (Malvaez et al., 2013; Stefanko et al., 2009). Habits are slow to form, being gradually acquired with repetition of successful actions in the presence of consistent stimuli, but once fully formed can be executed almost automatically, freeing attention to be focused elsewhere (Dickinson, 1985; Dolan and Dayan, 2013). The current data suggest that in the dorsal striatum the repressive enzyme HDAC3 could be a molecular ‘brake’ (McQuown and Wood, 2011; Vogel-Ciernia and Wood, 2012) on this type of learning, being engaged to slow habit formation and preserve behavioral control by the more cognitively-taxing, but less error-prone, goal-directed system until enough successful repetition has proceeded to ensure sufficient accuracy of the habit. In support of this, early in training, when habits do not yet dominate behavioral control, HDAC3 occupancy at the *Bdnf1* and *Nr4a2* promoters in the DLS and at the *Nr4a1* promoter in the DMS is enriched and then returns to baseline levels with the repeated training that promotes habit. That overexpression of HDAC3 in the dorsal striatum prevented development of behavioral habits further suggests that, under suitable conditions (e.g., repeated success), an instrumental learning opportunity triggers activity-dependent signaling that removes HDAC3 to create the transcriptionally-permissive state that allow habits to strengthen and, eventually, come to control behavior. Dorsal striatal HDAC3, therefore, normally curtails habit.

Consolidation of long-term memories, such as habits, depends on gene transcription (Alberini, 2009). HDAC3 represses transcription by removal of acetylation and by recruiting complementary repressive enzymes, such as methyltransferases or phosphatases. Removal of HDACs from gene promoters allows histone acetylation, which not only decreases affinity of histone tails for DNA, but also serves as a recruitment signal for transcriptional coactivators (Kouzarides, 2007) that can promote active gene expression subserving synaptic plasticity and learning (Borrelli et al., 2008; McQuown and Wood, 2011; Vogel-Ciernia and Wood, 2012). The data here provide evidence that both overtraining and HDAC inhibition disengage HDAC3 from specific gene promoters in both the DLS and DMS. Indeed, both *Bdnf1* and *Nr4a2* expression was induced in the DLS following intermediate training with HDAC inhibitor treatment. In several cases, however, gene expression did not perfectly follow HDAC3 occupancy at the promoter region. This could due to differential temporal dynamics between transcription of these genes and HDAC3 interactions with specific promoters (Kwapis et al., 2017). Moreover, HDAC3 does not function alone to regulate gene expression, rather a complex concert of factors, including additional chromatin modifying mechanisms, such as HATS (e.g., CREB-binding protein) and recruitment of other repressive mechanisms (Kennedy, 2013) are involved. Future experiments using genome-wide approaches may better elucidate the transcription underlying habit learning. Nevertheless, the data here reveal a crucial role for HDAC3 in the enhancing effects of HDAC inhibition on striatal-dependent habit memory.

Transcription regulated by CREB has been shown to be essential for long-term memory (Bourtchuladze et al., 1994; Guzowski and McGaugh, 1997; Pittenger et al., 2002; Yin et al., 1994), and, importantly, CREB function in the dorsal striatum is crucial for habit learning (Pittenger et al., 2006). The enhancement of memory by HDAC inhibition has been demonstrated to occur through regulation of CREB-regulated genes (McQuown et al., 2011; Vecsey et al., 2007), such as *Bdnf*, *Nr4a1*, and *Nr4a2* (Tao et al., 1998; Vecsey et al., 2007), genes themselves implicated in learning (Gottmann et al., 2009; Intlekofer et al., 2013; Lu et al., 2008; McNulty et al., 2012; McQuown et al., 2011; Vecsey et al., 2007; White et al., 2016). Here we found that as habits form HDAC3 was disengaged from the *Bdnf1* and *Nr4a2* promoters in the DLS and the *Nr4a1* promoter in the DMS and that HDAC inhibition, which accelerated habit formation, similarly altered HDAC3 occupancy at these promoters and also induced the expression of these genes. The data suggest that dorsal striatal HDAC3 might control habit formation via regulation of CREB-regulated genes.

Surprisingly, HDAC3 was found to negatively regulate habit in *both* the DLS and DMS. HDAC3 inhibition restricted to the DMS potentiated habit formation, whereas DMS-specific HDAC3 overexpression, while not affecting instrumental learning per se, prevented control from transitioning to dominance by the habit system, completely recapitulating the effect found with identical manipulations to the DLS. The overexpression result, along with data demonstrating HDAC3 enrichment early in training and removal with overtraining at the *Nr4a1* promoter in the DMS, provide evidence that DMS HDAC3 activity functions normally to repress habit. These results were unexpected in light of the canonical view that the DMS is crucial for action-outcome learning (Balleine et al., 2007; Balleine and O'Doherty, 2010) and evidence that DMS lesions force behavioral control by the habit system (Corbit and Janak, 2010; Yin et al., 2005a; Yin et al., 2005b). Previous data suggest that DLS circuits might store habit-related information (O'Hare et al., 2016; Rueda-Orozco and Robbe, 2015; Shan et al., 2015; Smith and Graybiel, 2013), with memories vital for goal-directed control stored in the DMS (Gremel et al., 2016; Gremel and Costa, 2013; Shan et al., 2014). One possibility is that HDAC3 may regulate the formation and storage of habit memories in the DLS, while in the DMS it depotentiates the action-outcome memories underlying goal-directed control as deliberation becomes less required and actions become chunked (Graybiel, 1998; Smith and Graybiel, 2016) into stereotyped units. Indeed, neurons in the DLS will show chunking-related activity that strengthens with training, while simultaneously DMS neurons show activity that wanes at deliberation points with extended training (Thorn et al., 2010). It is also possible that HDAC3 regulates the activity of DMS indirect-pathway projections, which have been shown to oppose goal-directed behavioral control and thereby promote habit (Li et al., 2016). In either case, the distinct HDAC3 occupancy patterns at learning-related gene promoters in the DLS v. DMS suggests differential transcriptional regulation by HDAC3 in these subregions. Importantly, these results challenge the strict dissociation between DMS and DLS function in goal-directed and habitual control of behavior.

These data reveal a new molecular mechanism regulating habit formation. HDAC3 functions as a molecular ‘gate’ over habit, remaining in place to slow the transition to habit and being removed when the conditions are ripe for habits to dominate. The balance between goal-directed and habitual control is disrupted in a variety of disease states, including addiction (Belin et al., 2013), alcoholism (Corbit and Janak, 2016), obesity (Corbit, 2016), obsessive-compulsive disorder (Gillan et al., 2011), autism-spectrum disorder (Alvares et al., 2016), schizophrenia (Morris et al., 2015), and Parkinson’s disease (Redgrave et al., 2010). An overreliance on habit is especially associated with the various forms of compulsivity that manifest across a range of psychiatric diseases (Gillan et al., 2016b; Voon et al., 2015). Moreover, stress, a predisposing condition to many psychiatric illnesses, may remove HDACs in abnormal ways (White and Wood, 2014), potentiating habits (Dias-Ferreira et al., 2009), to potentially promote maladaptive compulsive behavior. The current data, therefore, suggest a potential epigenetic mechanism for such maladaptive behavior and a target for therapeutic intervention.

## ACKNOWLEDGEMENTS

This research was supported by a Hellman Foundation Fellowship, a UCLA Faculty Career Development award, and grant DA035443 from NIDA, to KMW, grant DA025922 and AG051807 to MAW, and start-up funds from UCLA Life Sciences Division to PJK. We also thank Dr. Jeffrey Bye for statistical consulting.

## AUTHOR CONTRIBUTIONS

MM and KMW designed the research, analyzed and interpreted the data with assistance from MAW and PJK. MM conducted the research with assistance from VYG and MDM. DPM and MAW prepared and contributed viral constructs and conducted ChIP experiments. MM, NAA, and PJK collected and analyzed qRT-PCR and Western Blot data. MM and KMW wrote the manuscript with assistance from MAW and PJK.

## CONTACT FOR REAGENT AND RESOURCE SHARING

Further information and requests for resources and reagents should be directed to and will be fulfilled by the Lead Contact, Kate M. Wassum.

## EXPERIMENTAL MODEL AND SUBJECT DETAILS

Male, Long Evans rats (280-340 g at start of experiment; Charles River Laboratories, Wilmington, MA) were group housed and handled for 5-7 days prior to the onset of the experiment. Unless otherwise noted, separate groups of naïve rats were used for each experiment. Rats were food-restricted to ~85% free-feeding body weight with water provided *ad libitum* in the home cage. Experiments were performed during the dark phase of the 12:12 hr reverse dark/light cycle. All procedures were conducted in accordance with the NIH Guide for the Care and Use of Laboratory Animals and were approved by the UCLA Institutional Animal Care and Use Committee.

## METHOD DETAILS

### Surgery

For post-training microinfusions, rats were implanted with guide cannula (22-gauge stainless steel, Plastics One, Roanoke, VA) targeted bilaterally 1 mm above the DLS (AP: +1.0 mm; ML: ± 3.7; DV: -3.5) or DMS (AP: +1.0; ML: ± 2.0; DV: -3.5) using standard surgical procedures described previously (Malvaez et al., 2015). For viral vector infusions, two weeks prior to behavioral procedures, rats were randomly assigned to viral group, anesthetized using isoflurane and infused, as previously described (Malvaez et al., 2013; Malvaez et al., 2011), with adeno-associated virus expressing the HDAC3 point mutant, HDAC3Y298H (AAV2/1-CMV-HDAC3Y298H-V5), wild-type HDAC3 (AAV2/1-CMV-HDAC3-V5), or the empty vector (AAV2/1- CMV-EV). AAV2/1-HDAC3Y298H, AAV2/1-HDAC3, or AAV2/1-EV (0.50 μl) was infused at a rate of 6 μl/hr via an infusion needle positioned in the DLS (AP: +1.0 mm; ML: ± 3.7; DV: -4.6) or DMS (AP: +1.0; ML: ± 2.0; DV: -4.6). Coordinates were selected based on area of densest H4K8Ac following systemic HDAC inhibition. A nonsteroidal anti-inflammatory agent was administered pre- and post-operatively to minimize pain and discomfort. Following surgery, rats were individually housed and allowed to recover for 5-7 days (for cannula implants) or 2 weeks (for viral expression) prior to onset of behavioral training. Cannula placements were verified using standard histological procedures. Subjects were removed from the study if cannula were misplaced outside the DLS (*N*=3) or DMS (*N*=2). Restriction of virus to either the DLS or DMS was verified with immunofluorescence using an antibody to recognize the V5 tag. Subjects were removed from the study due to lack of expression or expression outside of the intended target zone (HDAC3Y98H: DLS, *N*=0, DMS, *N*=7; HDAC3: DLS, *N*=3, DMS, *N*=2).

### Apparatus

Training took place in a set of 16 Med Associates operant chambers (East Fairfield, VT) housed within sound- and light-attenuating boxes, described previously (Malvaez et al., 2015). The chambers contained a retractable lever to the left of a recessed food delivery port and a pellet dispenser that delivered a single 45-mg pellet (Bio-Serv, Frenchtown, NJ) when activated. A 3-watt, 24-volt house light mounted on the top of the wall opposite the food port provided illumination.

### Behavioral Procedures

#### Training

Each session began with the illumination of the houselight and insertion of the lever, where appropriate, and ended with the retraction of the lever and turning off of the houselight. Rats were given only one training session/day. All rats first received two sessions in which they were trained to retrieve food rewards (grain or chocolate pellets, counterbalanced across subjects) from the food-delivery port. Rewards were delivered on a random-time 60-s schedule and rats were given 30 total rewards/session. This was followed by daily instrumental training sessions in which food-pellet rewards could be earned by lever pressing. During the first instrumental training session, outcomes were delivered on a continuous reinforcement (CRF) schedule. After this, rats were was shifted to random-interval (RI)-30 s schedule of reinforcement for either 3 (limited training), 4 (intermediate training), or 6 (extended training) consecutive days. Each training session lasted until 30 outcomes had been earned or 40 min elapsed. In some cases, rats in the extended training condition were tested, as described below, then given an additional 6 consecutive days of instrumental training prior to being tested for a second time. For one experiment (Fig. S2), a random-ratio (RR) 10 schedule of reinforcement was substituted for the RI-30s schedule.

All rats were also given equated non-contingent access to the alternate food pellet (e.g., chocolate pellets if grain pellets served as the training outcome) in a different room and context (clear plexiglass cage) a minimum of either 2 hr before or after (alternated daily) each instrumental training session.

#### Devaluation test

Outcome devaluation testing was conducted after training. Rats were given 1 hr unlimited access to either the outcome previously earned by lever pressing (Devalued condition) or the previously exposed, alternate outcome to control for general satiety (Valued condition). Immediately after this pre-feeding, lever-press behavior was assessed during a brief, 5-min, non-reinforced probe test. Following the lever-pressing test, rats were given 10 min of exposure to each food pellet type, counterbalanced for order, in which the amount of each pellet type consumed was measured to ensure full rejection of the devaluated reward. Subjects who failed to consume less of the devalued v. non-devalued food type were not included in the analysis (Systemic NaBut intermediate group: Vehicle, *N*=1, NaBut, *N*=2; Systemic NaBut extended group: Vehicle, *N*=1, NaBut, *N*=1; Systemic NaBut 10 hr post training: Vehicle, *N*=1; DLS HDAC3: EV, *N*=1, HDAC3, *N*=3; DMS RGFP966: Vehicle, *N*=2, NaBut, *N*=1; DMS HDAC3Y298H: EV, *N*=6, HDAC3Y298H, *N*=3; DMS HDAC3: EV, *N*=4, HDAC3, *N*=2). Rats were given 2 days off, then 1 day of instrumental retraining (RI-30 s), prior to a second test in which they were pre-fed the opposite outcome, such that each rat was tested in both the Valued and Devalued conditions. The order of devaluation testing (Valued v. Devalued) was counterbalanced across subjects.

#### Omission test

Rather than the devaluation test, one group of subjects was given an omission test after training (Fig. S1). This consisted of a 30-min session in which the contingency between the lever press and the outcome was reversed. For this test, a food pellet was non-contingently delivered every 20 s. For rats in the omission condition, each lever press would reset the 20-s timer and thus delaying outcome delivery and would not, itself, earn reward. Each yoked-control subject received identical food pellet exposure to one rat in the omission group, with lever pressing having no programmed consequences.

### Delivery of histone deacetylase inhibitors

Sodium butyrate (NaBut; Sigma, St. Louis, MO) was dissolved in sterile water and delivered via intraperitoneal injection at a dose of 1000 mg/kg, 1.0 ml/kg immediately following each RI-30 s (or RR-10) instrumental training session, including the retraining session. This dose was selected based on its effectiveness in enhancing memory in a variety of other tasks (Bredy et al., 2007; Lattal et al., 2007; Malvaez et al., 2010; Ploense et al., 2013; Stefanko et al., 2009). RGFP966 (Abcam) was dissolved in DMSO and diluted in a vehicle of 30% hydroxypropyl-β-cyclodextrin and 100 mM sodium acetate, pH 5.4. The final DMSO concentration was 5% for drug and vehicle. RGFP966 (1.0 ng/0.5 μl/side over 1 min) was infused bilaterally immediately following each RI-30 s instrumental training session via injectors inserted into the guide cannula fabricated to protrude 1 mm ventral to the cannula tip using a microinfusion pump. For drug group assignment, subjects were counterbalanced based on lever-press rate during the CRF instrumental training session.

### Viruses

Wild-type HDAC3 was amplified from mouse hippocampal cDNA and cloned into a modified pAAV-IRES-hrGFP (Agilent), under control of the CMV promoter and Beta-globin intron. The 3x-FLAG tag, IRES element, and hrGFP were removed from the vector and replaced with a V5 tag, allowing for a c-terminal fusion to HDAC3 (MW91). To create the point mutation, a single nucleotide substitution in exon 11 to direct production of a histidine residue in place of tyrosine at amino acid 298 was created (HDAC3Y298H; MW92). For the empty vector control, the HDAC3 coding sequence was not present, but all other elements remained (MW87). Adeno-associated virus (AAV) was made by the Penn Vector Core (University of Pennsylvania) from the above described plasmids and was serotyped with AAV 2.1.

### Immunofluorescence

To assess histone acetylation induced by post-training HDAC inhibitor treatment, a separate group of rats were perfused 1 hr following the last session of intermediate training session and either vehicle or NaBut injection. Rats that received post-training microinfusions, after all training and testing, were given a final training session followed by either vehicle or RGFP966 treatment 1 hr prior to perfusion. Rats that received viral manipulations were taken from their home cages and perfused at the conclusion of behavioral testing. Rats were transcardially perfused with PBS followed by 10% formalin. The brains were removed, post-fixed in formalin, then cryoprotected, cut with a cryostat at a thickness of 30 μm and collected in PBS. Immunohistochemical analysis was performed as described previously (Malvaez et al., 2013; Malvaez et al., 2011; Malvaez et al., 2010). Briefly, floating coronal sections were blocked for 1 hr at room temperature in 8% normal goat serum (NGS, Jackson ImmunoResearch Laboratories) with 0.3% Triton X-100 in PBS and then incubated overnight at 4°C in 2% NGS, 0.3% Triton X-100 in PBS with primary antibody (anti-H4K8Ac, 1:1000, Cell Signaling, cat. no. 2594; anti-V5, 1:1000, Abcam, cat. no. 9116). The sections were then incubated for 2 hr at room temperature with goat anti-rabbit IgG-FITC (1:1000, Invitrogen, cat. no. A11070). All sections were washed 3 times for 5 min each in PBS before and after each incubation step and mounted on slides using ProLong Gold antifade reagent with DAPI (Invitrogen).

All images were acquired using a Keyence (BZ-X710) microscope with a 20X objective (CFI Plan Apo), CCD camera, and BZ-X Analyze software. A single optimized acquisition exposure time was used for all images acquired from a particular slide with all treatment groups represented. Immunolabeling was quantified using ImageJ software (NIH) by measuring the optical density in the area of interest from comparable 20X images. For each animal, the average of three sections was calculated to give a mean optical density.

### RNA isolation and qPCR

Tissue was collected 1 hr following the last training session/injection by taking 1 mm sections and microdissecting the DLS and DMS on cold ice. Frozen DLS and DMS punches were homogenized in TRIzol and processed according to the manufacturer's instructions. RNA was purified with RNeasy Micro columns (QIAGEN) and spectroscopy confirmed that the RNA had 260/280 and 260/230 ratios >1.8. RNA was reversed transcribed into cDNA using iScript cDNA synthesis (Bio-Rad). Gene expression analysis was performed using Fluidigm DELTAgene^TM^ real-time qPCR assays. Each reaction was analyzed according to the standard ΔΔ*C*t method using glyceraldehyde-3-phosphate dehydrogenase (GAPDH) as a normalization control. Samples were then normalized to vehicle home cage controls.

Primers used for real-time PCR were as follows: GAPDH: F-GTGGACCTCATGGCCTACAT R-TGTGAGGGAGATGCTCAGTG; BDNF1: F-CAGGAGCGTGACAACAATGTGA, RACCATAGTAAGGAAAAGGATGGTCAT; Nr4a1: F-CCGGTGACGTGCAGCAATTTTATGAC, R-GGCTAGAATGTTGTCTATCCAGTC; Nr4a2: F-ATTGCTGCCCTGGCTATGGT, RGACCATCCCATTATTGAAAGTCACATGGTC; c-Fos: F-GGACAGCCTTTCCTACTACCA, RCGGACAGATCTGCGCAAAA; FosB: F-AGAGCCAGGCCTAGAAGACC, R-TTGTTCCGCTCTCTGCGAAC; Egr1: F-AACCCTACGAGCACCTGAC, R-CGGGTAGTTTGGCTGGGATA.

### Chromatin immunoprecipitation

ChIP was performed as described previously (Malvaez et al., 2013), based on the protocol from the Magna ChIP kit (cat# 17-610, Millipore). Tissue was collected 1 hr following the last training session/injection. Brains were rapidly extracted on cold ice and sliced into 1-mm sections. For each sample, bilateral DLS and DMS tissue was isolated and rapidly frozen. Chromatin was immunoprecipitated overnight with anti-HDAC3 or nonimmune IgG (cat# 17-10238, Millipore) and Magna ChIP Protein A+G Magnetic Beads (cat# 16-663, Millipore), DNA promoter enrichment was quantified using quantitative PCR using the Roche 480 Lightcycler and SYBR Green (cat# 04707516001, Roche). To normalize ChIP-qPCR data, the percent input method was used. The input sample was adjusted to 100% and both the IP and IgG samples were calculated as a percent of this input using the formula: 100*AE^(adjusted input - Ct(IP)). An in-plate standard curve determined amplification efficiency (AE). Fold enrichment was calculated as a ratio of the ChIP to the average IgG and samples were then normalized to vehicle-treated home cage controls. Primers used for ChIP were as follows: Bdnf Exon I: F-ACGTCCGCTGGAGACCCTTAGT RGGCAGCCTCTCTGAGCCAGTTA; Nr4a1: F-ACCACCTTCACATCCCTCAG R-CAGCAGCTCAGTCAAGTCAC; Nr4a2: F-ATCTGTCCAGCCAAGTCTCG R-GGGGAACTCGGGAAAGGTAA.

## QUANTIFICATION AND STATISTICAL ANALYSIS

### Behavioral analysis

Lever presses were the primary behavioral output measure and were collected continuously for each training and test session. To assess acquisition of the instrumental task, lever-press rate was averaged across each instrumental training session and plotted over training days. For the devaluation test we normalized lever pressing for the Valued or Devalued states over total lever pressing (Valued + Devalued state pressing), as described previously (Gremel et al., 2016; Gremel and Costa, 2013).

### Statistical analysis

Datasets were analyzed by two-sided Student’s *t* tests, one- or two-way repeated-measures analysis of variance (ANOVA), as appropriate. Bonferroni or Dunnet’s corrected *post hoc* tests were performed to clarify all main effects and interactions. Two-tailed, paired t-tests were used for *a priori* planned comparisons, as advised by (Levin et al., 1994) based on a logical extension of Fisher’s protected least significant difference (PLSD) procedure for controlling familywise Type I error rates. All datasets met equal covariance assumptions as analyzed using Levene’s and Box’s M tests, justifying ANOVA interpretation (Tabachnick et al., 2001). Alpha levels were set at *P*<0.05.

## DATA AVAILABILITY

All data that support the findings of this study are available from the corresponding author.

